# Checklist and practical identification key for the cichlid fishes (Cichliformes: Cichlidae) of the La Plata drainage in Bolivia, including three new geographical records

**DOI:** 10.1101/785006

**Authors:** Pascal I. Hablützel, Robert B. Huanto

## Abstract

In comparison with the Bolivian Amazon, the ichthyofauna of the La Plata drainage of Bolivia received relatively little attention historically. Until now, 14 species of cichlid fish have been registered from this area. After an exhaustive review of museum collections (Museo de Historia Natural Noel Kempff Mercado y Colección Boliviana de Fauna), we can report three additional species: *Astronotus crassipinnis* (Heckel, 1840), *Mesonauta festivus* (Heckel, 1840) and *Satanoperca pappaterra* (Heckel, 1840). Four other species, which have been listed in previous publications, can be confirmed for the La Plata drainage of Bolivia based on the examination of voucher specimens: *Aequidens plagiozonatus* Kullander, 1984, *Apistogramma commbrae* (Regan, 1906), *A. trifasciata* (Eigenmann & Kennedy, 1903) and *Crenicichla vittata* Heckel, 1840. As such, 16 of the 17 species can be referenced with voucher specimens in museum collections. We also provide an identification key for the cichlid fish species of the study area.

## Resumen

En comparación con la Amazonía boliviana, la ictiofauna de la Cuenca de la Plata en Bolivia recibió relativamente poca atención históricamente. Hasta el momento, se han registrado 14 especies de peces cíclidos de esta área. Después de una revisión exhaustiva de las colecciones de los museos (Museo de Historia Natural Noel Kempff Mercado y Colección Boliviana de Fauna), podemos informar que se adicionan tres especies: *Astronotus crassipinnis* (Heckel, 1840), *Mesonauta festivus* (Heckel, 1840) y *Satanoperca pappaterra* (Heckel, 1840). Otras cuatro especies, que han sido incluidas en publicaciones anteriores, pueden confirmarse para la Cuenca de La Plata en Bolivia con base en revisiones de especímenes *voucher*: *Aequidens plagiozonatus* Kullander, 1984, *Apistogramma commbrae* (Regan, 1906), *A. trifasciata* (Eigenmann & Kennedy, 1903) y *Crenicichla vittata* Heckel, 1840. Como tal, 16 de las 17 especies pueden ser referenciadas con especímenes de lotes en colecciones de los museos. También proporcionamos una clave de identificación para las especies de cíclidos del área de estudio

Palabras clave

Biogeografía, Peces de agua dulce, río Paraguay, nuevo reporte, Lista de verificación

## Introduction

The fish fauna of Bolivia is one of the richest in the world (Sarmiento & Barrera 2004; Carvajal-Vallejos *et al.* 2014; Sarmiento *et al.* 2014). Unfortunately, it remains poorly documented in the scientific literature. This is particularly the case for the Bolivian part of the La Plata basin, which comprises the Paraguay river in the eastern part of the country, and the rivers Pilcomayo and Bermejo in the south (Figure 1). The first accounts on fish in the Bolivian part of the La Plata basin that include cichlids have been made by Haseman in 1909, who collected fish for the Carnegie Museum (USA). All cichlids he reported from Bolivia are from Puerto Suárez (Haseman 1911) in the río Paraguay drainage. The location ‘São Francisco, Bolivia’ in Haseman’s (1911) work is actually located in Brazil (‘Rio São Francisco, into Paraguay river, Matto Grosso’; Eigenmann (1911)). In the 1950’s, Travassos collected several fish species, including cichlids, in the Paraguay and Otuquis drainages, but did not identify them at species level (Travassos 1957). In 1970, Terrazas Urquidi published a first list of the Bolivian fish species. The sources for his reports on *Apistogramma taeniata*, *A. trifasciata*, *A. corumbae* and *Crenicichla vittata* from the Bolivian río Paraguay drainage remain, however, unclear. In the 1980s, Kullander wrote a series of taxonomic accounts on the cichlid fishes of the río Paraguay drainage (Kullander 1981, 1982c, b, a, 1984, 1987). However, with the exception of the first descriptions of *A. inconspicua* and *Bujurquina oenolaemus*, he did not include Bolivian specimens.

**Figure 1:**
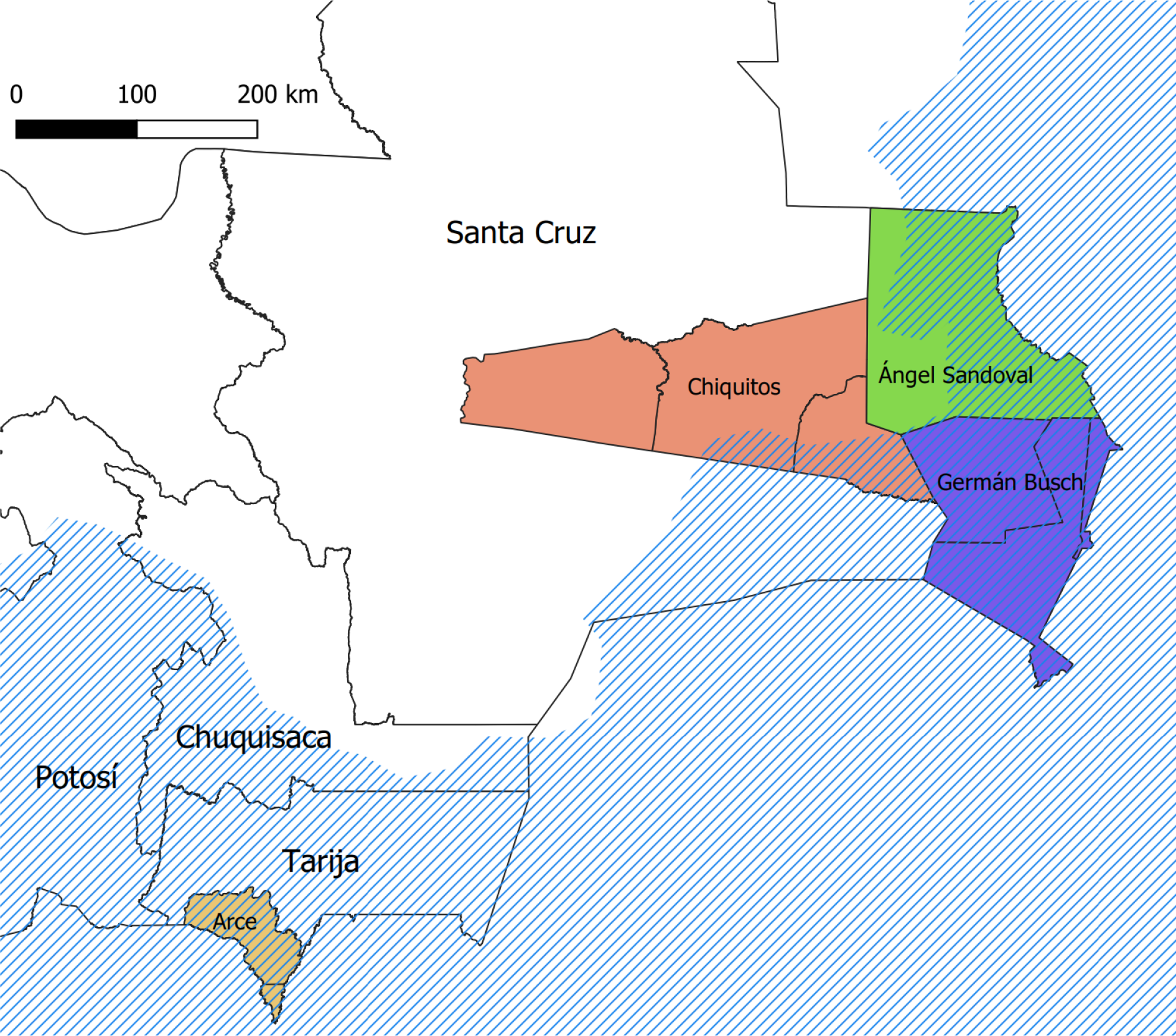
Map of the study area. The blue shaded area indicates the La Plata drainage, covering the Tarija department and parts of Santa Cruz, Chuquisaca and Potosí. Lots in museum collections have been reported to be from the provinces Arce (Tarija), Chiquitos, Ángel Sandoval and Germán Busch (all Santa Cruz).

Since the 1990’s several extensive surveys have been conducted by Bolivian researchers mostly affiliated to the Museo de Historia Natural ‘Noel Kempff Mercado’ (MHNNKM) in Santa Cruz de la Sierra. With the notable exception of the article by Farell & Cancino (2007), these studies have been published in governmental reports and academic theses which are unfortunately not readily accessible. For most of the species lists published in governmental reports, little metadata (such as sampling sites, dates, information on species identification, pictures, or lists of voucher specimens etc.) is available. Because of these limitations and since the species identification is not verifiable in most cases, we refrain from citing this grey literature in the present study, although we consider its content very valuable.

Geographical species records provide the basis of local, regional or national inventories on which biodiversity measures can be estimated, they can be used to study biogeographical patterns or they enable us to monitor changes in biodiversity through time. Regarding this importance, the publication of such records should follow certain standards allowing the verification of species identification. In this study, we report on the species of cichlid fishes from the Bolivian part of the La Plata drainage. We aim not only to report a species list, but also to link records to voucher specimens in museum collections, to provide photographs of new records and to facilitate future identification with an identification key.

## Study area

The study area includes the Bolivian part of the La Plata drainage in the departments Santa Cruz, Chuquisaca and Tarija. The major sub-drainages in this area are the rivers Paraguay, Pilcomajo and Bermejo.

## Methods

We report the cichlid species present in the collections of the Colección Boliviana de Fauna and the Museo de Historia Natural ‘Noel Kempff Mercado’ in La Paz and Santa Cruz de la Sierra, respectively. Fish re-identification took place in 2010. Besides a few exceptions, newer collections are therefore not considered in this study. The list of lots in this study should not be considered complete for various reasons (e.g. not being available at the time the examination took place). Fish were identified using the primary taxonomic literature (see references). Many collection numbers in the MHNNKM have been changed in recent years. To avoid confusion, we also provide the old numbers in brackets. Abbreviations: MNKP = Museo de Historia Natural ‘Noel Kempff Mercado’ (P = peces). CBF = Colección Boliviana de Fauna. To assess the completeness of the species list provided in this study statistically, we estimated rarefaction curves using the rarecurve function of the package vegan v2.5-2 (Oksanen *et al.* 2007) in R v3.4.2 (R Development Core Team 2012).

## Results

### *Aequidens plagiozonatus* Kullander, 1984

*Aequidens plagiozonatus* Kullander, 1984

#### Type locality

Brazil, State of Mato Grosso, R. Paraguay system Mun. Itiquira, internal lakes of the Piquiri-Itiquira system, Fazenda Santo Antonio do Paraíso.

##### Bibliography for the study area

*Aequidens plagiozonatus*; Farell & Cancino (2007): localities (Río Aguas Calientes).

###### Examined material

**Bolivia: Santa Cruz: Angel Sandoval:** MNKP 4765 (3518), 4, Curiche Tapera, K. Osinaga & P. Coro, 29 Sep 1999.

###### Identification

Species most frequently misidentified as members of the genus *Aequidens* belong to genera *Bujurquina*, *Cichlasoma* and *Laetacara*. *Bujurquina* differs from the other genera by having conspicuous lateral and nape bands. *Laetacara* has a dark stripe from the eye to the tip of the snout and a scaled preoperculum (vs. no dark stripe and preoperculum naked). *Cichlasoma* differs from the other genera in having scale rows in the proximal area of the rayed part of the dorsal fin and 6 vertical bars behind the midlateral spot (vs. 5 in *Aequidens*). *Aequidens plagiozonatus* is the only species of its genus to have been reported and confirmed from the upper río Paraguay drainage. It is medium sized and reaches 10.3 cm SL (Kullander 1984). See Figure 2 for general appearance. A detailed description and diagnosis can be found in Kullander (1984).

**Fig. 2:**
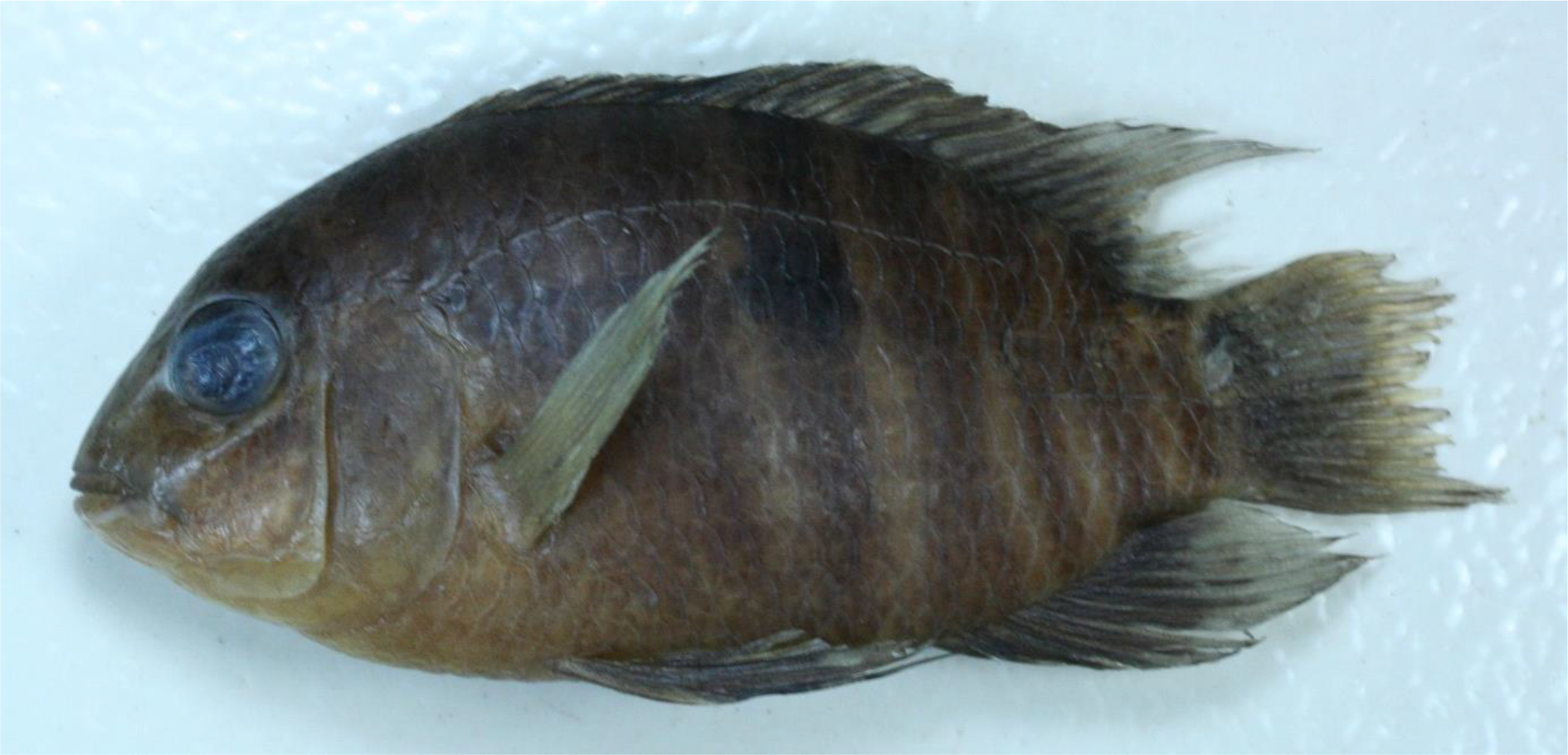
*Aequidens plagiozonatus* from the curiche Tapera, río Paraguay drainage, Angel Sandoval, Santa Cruz, Bolivia (MNKP 6598 (3518)).

### *Apistogramma borellii* (Regan, 1906)

*Heterogramma borellii* Regan, 1906

*Heterogramma ritense* Haseman, 1911

*Heterogramma rondoni* Miranda Ribeiro, 1918

*Apistogramma aequipinnis* Ahl, 1938

*Apistogramma reitzigi* Mitsch, 1938

#### Type locality

Carandasiñho, Matto Grosso.

##### Bibliography for the study area

*Heterogramma borellii*; Haseman (1911): locality (Puerto Suarez) and voucher material.

*Apistogramma borellii*; Terrazas Urquidi (1970): listing (R. Paraguay).

*Apistogramma borellii*; Schindler & Staeck (1993): listing.

###### Examined material

**Bolivia: río Paraguay drainage: Santa Cruz: Angel Sandoval:** MNKP 3215 (3927), 1, Laguna Mandioré, K. Osinaga, 15 Jul 1998. MNKP 4586 (3504), 12, Hacienda Vista Hermosa, laguna, K. Osinaga & P. Coro, 19 Sep 1999. **Germán Busch:** MNKP 1177 (1643), 4, Puerto Suárez, Laguna Cáceres, San Eugenio, K. Osinaga, R. Coca, V. Fuentes & A. Franco, 19 Feb 1996. MNKP 1178 (1646), 1, Puerto Suárez, Laguna Cáceres, K. Osinaga, R. Coca, V. Fuentes & A. Franco, 22 Feb 1996. MNKP 1180 (1667), 2, Laguna Cáceres, K. Osinaga, R. Coca, V. Fuentes & A. Franco, 22 Feb 1996 (lot also contains 1 specimen of *Cichlasoma dimerus*). MNKP 1183 (1671), 3, Puerto Suárez, Laguna Cáceres, Palo Santo, K. Osinaga, R. Coca, V. Fuentes & A. Franco, 22 Feb 1996. MNKP 1217 (2260), 4, Puerto Quijarro, Tamarinero, Canal Tamengo, K. Osinaga, R. Coca, V. Fuentes & A. Franco, 26 Feb 1996. MNKP 1219 (2732), 10, Puerto Quijarro, bahia Cáceres, Canal Tamengo, K. Osinaga, R. Coca, V. Fuentes & A. Franco, 26 Feb 1996 (lot also contains 1 specimen of *Cichlasoma dimerus*). MNKP 1222 (4229), 5, Puerto Suarez, Laguna Cáceres, curiche temporal, K. Osinaga, R. Coca, V. Fuentes & A. Franco, 19 Feb 1996. MNKP 1393 (1491), 2, Puerto Suárez, Laguan Cáceres, P. Rebolledo, K. Osinaga, V. Fuentes, L. Paniagua & L. Paredes, 9 Aug 1996. MNKP 1401 (1496), 1, Puerto Suárez, río San Lorenzo, P. Rebolledo, K. Osinaga, V. Fuentes, L. Paniagua & L. Paredes, 6 Aug 1996. MNKP 1416 (1517), 3, Puerto Suárez, Laguna Cáceres, P. Rebolledo, K. Osinaga, V. Fuentes, L. Paniagua & L. Paredes, 6 Aug 1996. MNKP 1444 (1560), 2, Puerto Quijarro, Laguna Cáceres, P. Rebolledo, K. Osinaga, V. Fuentes, L. Paniagua & L. Paredes, 13 Aug 1996. MNKP 1445 (1566), 11, Puerto Suarez, Laguna Cáceres, P. Rebolledo, K. Osinaga, V. Fuentes, L. Paniagua & L. Paredes, 9 Aug 1996. MNKP 1511 (2255), 18, Puerto Quijarro, Tamarinero, Laguna Cáceres, P. Rebolledo, K. Osinaga, L. Paniagua, L. Paredes & V. Fuentes, 13 Aug 1996. MNKP 1518 (2734), 5, Puerto Suárez, Laguna Cáceres, P. Rebolledo, K. Osinaga, L. Paniagua, L. Paredes & V. Fuentes, 11 Aug 1996. MNKP 1520 (2743), 1, Puerto Quijarro, Tamarinero, Laguan Cáceres, P. Rebolledo, K. Osinaga, L. Paniagua, L. Paredes & V. Fuentes, 15 Aug 1996. MNKP 2402 (2727), 2, Puerto Suarez, Laguna Cáceres, M.A. Parada & R. Yañez, 25 May 1997. MNKP 3882 (3055), 7, Puerto Suarez, Laguna Cáceres, K. Osinaga & J. Cardona, 26 Oct 1998. MNKP 3922 (4064), 6, río Pimiento, Laguan Cáceres, K. Osinaga & J. Cardona, 28 Oct 1998. MNKP 3932 (4175), 2, Laguna Cáceres, curichi Estancia Verdem, K. Osinaga & J. Cardona, 28 Oct 1998. MNKP 3954 (3219), 59, poza sobre el camino a la hacienda Santa Elena (Otuquis), K. Osinaga & J. Cardona, 31 Oct 1998 (lot also contains 3 specimens of *A. commbrae*). MNKP 3963 (6153), 4, poza sobre el camino a la hacienda Santa Elena (Otuquis), K. Osinaga & J. Cardona, 31 Oct 1998. MNKP 4010 (3886), 10, Bahia Corea, ‘Otuquis’, K. Osinaga, J. Cardona, A. Justiniano & M. Chavez, 5 Nov 1998. MNKP 4041 (3054), 1, El Carmen, Est. Campo en medio camino a 2 km al suroeste, K. Osinaga & J. Cardona, 10 Nov 1998.

###### Identification

To date, four species of *Apistogramma* have been reported from the upper río Paraguay drainage: *A. borellii*, *A. commbrae*, *A. inconspicua* and *A. trifasciata* (reports of other species exist, but should be considered questionable). *Apistogramma borellii* differs from *A. trifasciata* by the absence of an oblique black stripe between pectoral and anal fin origin and from *A. commbrae* and *A. inconspicua* by having the posteriormost vertical bar and caudal spot separated. Preserved specimen show a series of separated midlateral spots on the flanks. *Apistogramma borellii* is a small species reaching a size of 3.9 cm SL (Kullander 2003).

### *Apistogramma commbrae* (Regan, 1906)

*Heterogramma commbrae* Regan, 1906

*Heterogramma corumbae* Eigenmann & Ward, 1907

#### Type locality

Carandasiñho, Matto Grosso; Colonia Risso.

##### Bibliography for the study area

*Apistogramma commbrae*; Farell & Cancino (2007): locality (Río Tucavaca).

*Apistogramma corumbae*; Terrazas Urquidi (1970): listing (R. Paraguay).

###### Examined material

**Bolivia: río Paraguay drainage: Santa Cruz: Angel Sandoval:** MNKP 3216 (3927), 1, Laguna Mandioré, K. Osinaga, 15 Jul 1998. MNKP 4591 (3504), 1, Hacienda Vista Hermosa, laguna, K. Osinaga & P. Coco, 20 Sep 1999. **Chiquitos:** MNKP 5270, 13, Candelaria, río Tucavaca, M.E. Farell, M.Á. Velásquez & P. Hinojosa, 29 Dec 2003. **Germán Busch:** MNKP 1488 (1526), 5, Puerto Suarez, Laguna Cáceres, P. Rebolledo, K. Osinaga, V. Fuentes, L. Paniagua & L. Paredes, 13 Aug 1996. MNKP 2949 (1966), 1, Puerto Suarez, Hacienda Arcoiris, Laguna Cáceres, M.A. Parada & V. Fuentes, 1997. MNKP 3954 (3219), 3, poza sobre el camino a la hacienda Santa Elena, K. Osinaga & J. Cardona, 31 Oct 1998 (lot also contains 59 specimens of *C. borellii*). MNKP 3981 (3236), 1, poza camino a Santa Elena a 10 km de hacienda Las Camelias, K. Osinaga & J. Cardona, 1 Nov 1998. MNKP 6599 (3522), 1, río Santo Rosario, K. Osinaga & P. Coro, 1 Oct 1999.

###### Identification

*Apistogramma commbrae* differs from *A. trifasciata* by the absence of an oblique black stripe between pectoral and anal fin origin and from *A. borellii* by having the posteriormost vertical bar and caudal spot fused to a tail spot. In *A. commbrae*, the dark midlateral band is clearly fused with the tail spot, while in *A. inconspicua*, the band is somewhat separated from it. Preserved specimen of *A. commbrae* show several typically conspicuous longitudinal stripes along the flanks below the lateral band (less conspicuous or absent in *A. inconspicua*). Additional characters to distinguish *A. commbrae* from *A. inconspicua* are: modally 16 dorsal fin spines (vs. 15) and teeth on the posteromedial portion of the pharyngeal tooth plate bicuspid (vs. tricuspid; Kullander, 1982c). It is a small species reaching 32.6 mm SL (Kullander 1982b). See Figure 3 for general appearance. A detailed redescription of *A. commbrae* can be found in Kullander (1982b).

**Fig. 3:**
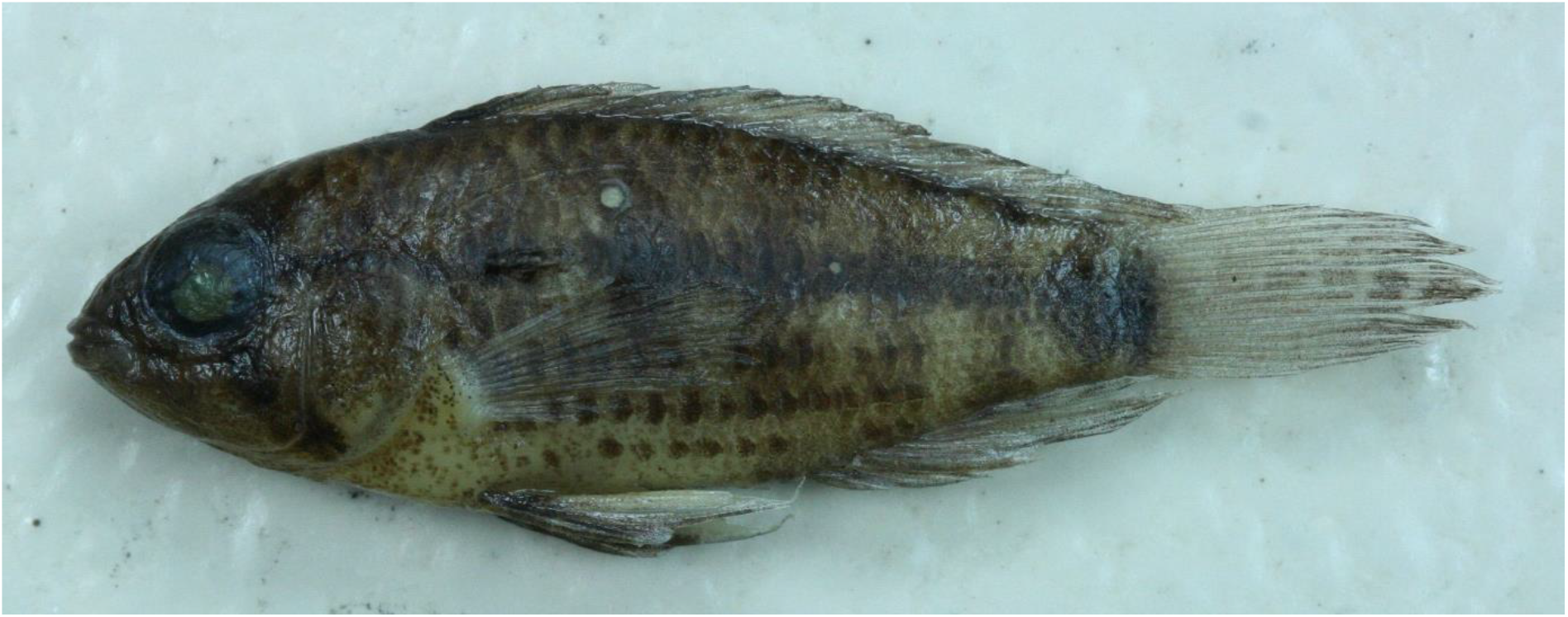
*Apistogramma commbrae* (female) from the río Tucavaca at Candelaria, Chiquitos, Santa Cruz, Bolivia (MNKP 5270).

### *Apistogramma trifasciata* (Eigenmann & Kennedy, 1903)

*Biotodoma trifasciatus* Eigenmann & Kennedy, 1903

*Heterogramma trifasciatum masiliense* Haseman, 1911

*Apistogramma trifasciatum haraldschultzi* Meinken, 1960

#### Type locality

Arroyo Chagalalina [Paraguay].

##### Bibliography for the study area

*Apistogramma trifasciata*; Terrazas Urquidi (1970): listing (R. Paraguay).

*Apistogramma trifasciata*; Schindler & Staeck (1993): listing.

###### Examined material

**Bolivia: río Paraguay drainage: Santa Cruz: Angel Sandoval:** MNKP 3188 (3921), 7, Puerto Gonzalo, río Pando, K. Osinaga, 12 Jul 1998. MNKP 3214 (3927), 2, Laguna Mandioré, K. Osinaga, 15 Jul 1998. MNKP 4592 (3504), 1, Hacienda Vista Hermosa, laguna, K. Osinaga & P. Coro, 20 Sep 1999.

###### Identification

Differs from all other species of *Apistogramma* by having an oblique black stripe between pectoral and anal fin origin. This stripe is absent in living specimens of at least one population in the Bolivian Amazon (personal observation), but seems to be always present in preserved animals. *Apistogramma trifasciata* is a small species reaching a size of 3.8 cm SL (Kullander 2003). See Figure 4 for general appearance of the species.

**Fig. 4:**
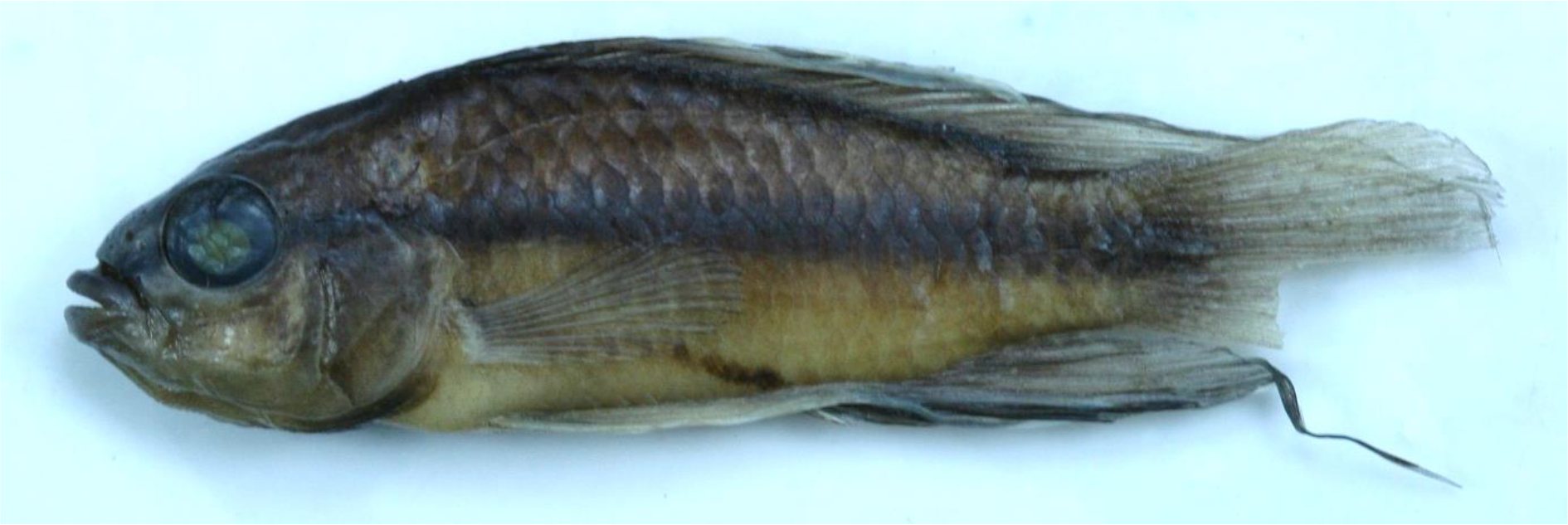
*Apistogramma trifasciata* (male) from the río Paragauy drainage in Bolivia (CBF 6321; exact locality unknown).

### *Apistogramma* sp. (unidentified juveniles)

#### Examined material

**Bolivia: río Paraguay drainage: Santa Cruz: Ángel Sandoval:** MNKP 4611 (3453), 6, laguna, Hacienda Vista Hermosa, laguna, K. Osinaga & P. Coro, 21 Sep 1999.

### *Astronotus crassipinnis* (Heckel, 1840)

*Acara crassipinnis* Heckel, 1840

#### Type locality

Rio-Paraguay … in Buchten … bei Villa Maria und Caiçara … Rio-Guaporè bei Matogrosso, im Rio-negro und im Rio-branco.

##### Bibliography for the study area

###### Examined material

**Bolivia: río Paraguay drainage: Santa Cruz: Ángel Sandoval:** MNKP 4810 (3600), 1, Santo Rosario, río Santo Rosario, K. Osinaga & P. Coro, 1 Oct 1999. **Germán Busch:** MNKP 1262 (1801), 1, Puerto Suarez, Hacienda Arcoiris, Laguna Cáceres, K. Osinaga, R. Coca, V. Fuentes & A. Franco, 28 Feb 1996. MNKP 1430 (1532), 1, Puerto Suarez, Castrillo, Laguna Cáceres, P. Rebolledo, K. Osinaga, V. Fuentes, L. Paniagua & L. Paredes, 9 Aug 1996. MNKP 3874 (3796), 1, Puerto Suarez, río Pimiento, K. Osinaga & J. Cardona, 23 Oct 1998.

###### Identification

*Astronotus crassipinnis* can been identified on the basis of the following unique combination of characters: ‘African type’ lips (see Figure 12 in Kullander (1986)), unpaired fins densely scaled, about 35 scales in the longitudinal row, no ocelli along the dorsal-fin base. This species is the only one of its genus known to occur in the Río Paraguay basin. It is a large species reaching 24 cm SL (Kullander 2003). See Figure 5 for general appearance. More detailed accounts on the appearance of *A. crassipinnis* are provided by Kullander (1981, 1986).

**Fig. 5:**
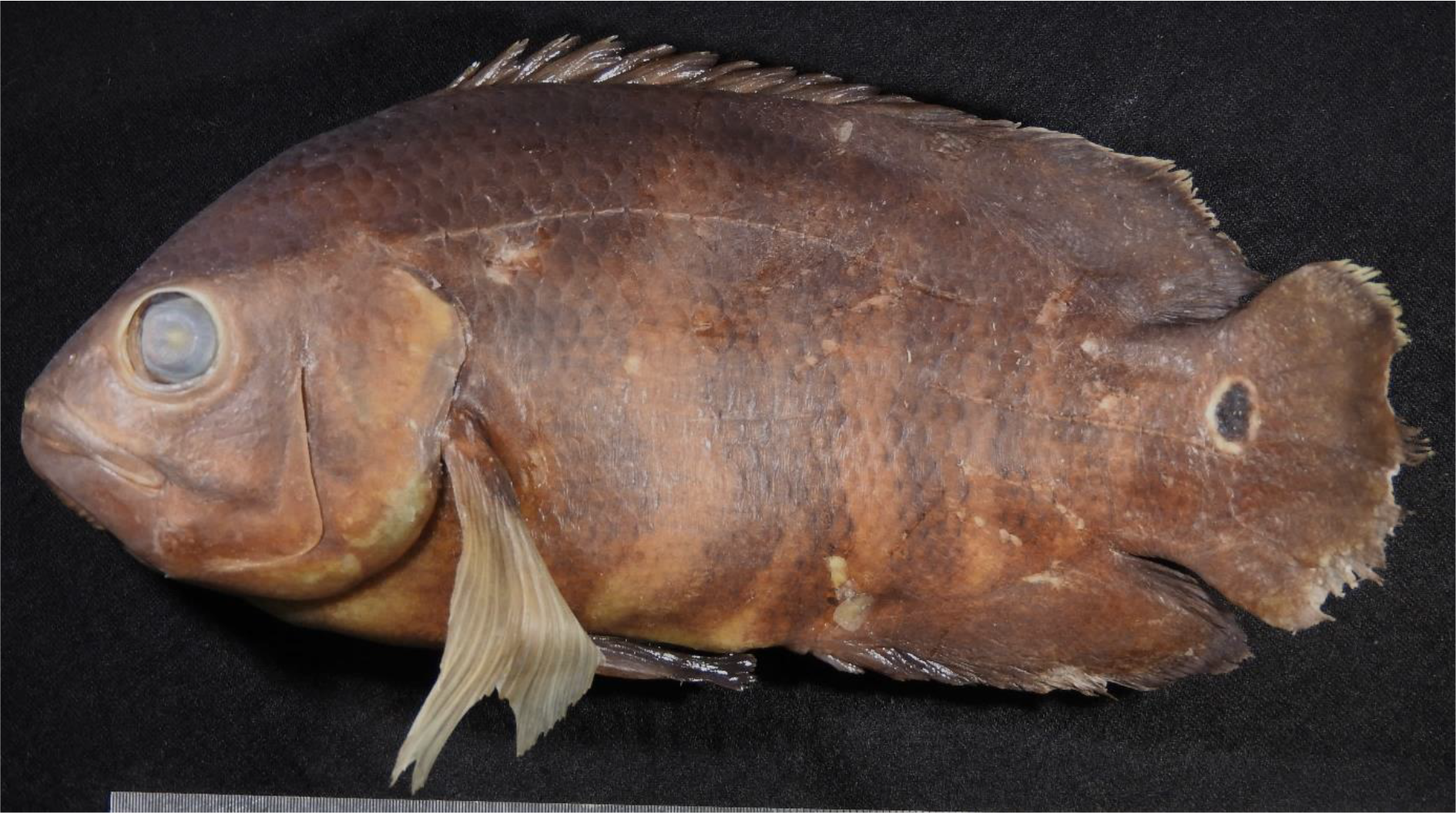
*Astronotus crassipinnis* from río Santo Rosaria at Santo Rosario, río Paraguay drainage, Angel Sandoval, Santa Cruz, Bolivia (MNKP 4810).

### *Bujurquina oenolaemus* Kullander 1987

*Bujurquina oenolaemus* Kullander, 1987

#### Type locality

Bolivie, dép. Santa Cruz. Rio Aguas Calientes à Aguas Calientes, à 25 km à l’est de Roboré, sur le rail. (Basin du Paraguay).

##### Bibliography for the study area

*Bujurquina oenolaemus*; Kullander (1987): diagnosis, description, photograph, illustration, locality (Río Aguas Calientes) and voucher specimen.

*Bujurquina oenolaemus*; Farell & Cancino (2007): localities (Río Aguas Calientes) and photography.

*Bujurquina oenolaemus*; Sarmiento *et al.* (2014): listing (Río Aguas Calientes).

###### Examined material

**Bolivia: río Paraguay drainage: Santa Cruz: Chiquitos:** MNKP 4692 (3481), 11, Aguas Calientes, río Aguas Calientes, K. Osinaga & P. Coro, 26-27 Sep 1999. MNKP 5202 (4142), 3, Aguas Calientes, río Aguas Calientes, V. Fuentes & S. Sangueza, 13 Dec 2000. MNKP 5244, 1, Localidad Aguas Calientes, El Playon, naciente del río Aguas Calientes, M.E. Farell, M.Á. Velásquez & P. Hinojosa, 5 Jan 2004. MNKP 5282, 6, punto de union de los ríos Jesus y Aguas Calientes, localidad Aguas Calientes, M.E. Farell, M.Á. Velásquez & P. Hinojosa, 4 Jan 2004. MNKP 5298, 9, Aguas Calientes, balneario El Burriño, río Aguas Calientes, M.E. Farell, M.Á. Velásquez & G. Huanca, 5 Jul 2004. MNKP 5352, 1, Aguas Calientes, zona El Playón, río Aguas Calientes, M.E. Farell, M.Á. Velásquez & G. Huanca, 5 Jul 2004. MNKP 5362, 2, Aguas Calientes, punto de union río Aguas Calientes y río Jesus, M.E. Farell, M.Á. Velásquez & G. Huanca, 5 Jul 2004.

###### Identification

Differs from *B. vittata* by having a shorter pectoral fin (29-33 vs. 36-41 % of SL) and a longer head (38-41 vs. 35-38 % of SL). Body with a dark reddish/ wine-coloured hue when alive, brachiostegal membrane reddish. *Bujurquina oenolaemus* is a small species reaching 7.1 cm SL (Hablützel, unpublished data).

### *Bujurquina vittata* (Heckel, 1840)

*Acara vittatus* Heckel, 1840

*Aequidens paraguayensis* Eigenmann & Kennedy, 1903

#### Type locality

Sümpfen um Cujabá, der Hauptstadt in der Provinz Matagrosso [sic].

##### Bibliography for the study area

*Aequidens paraguayensis*; Haseman (1911): localities (Puerto Suarez) and voucher material.

*Aequidens paraguayensis*; Terrazas Urquidi (1970): listing (Río. Pilcomayo and Río. Paraguay).

*Bujurquina vittata*; Farell & Cancino (2007): localities (Río Otuquis drainage).

###### Examined material

**Bolivia: río Paraguay drainage: Santa Cruz: Angel Sandoval:** CBF 8679, 1, Pantanal, Río Las Conchas, Estancia Las Conchas, arroyo “submontano” de “aguas negras”, con presencia de grandes pozas, rápidos y cataratas, 17° 33’ 58.3’’S 59° 28’ 17.1’’W, J. Sarmiento & K. Osinaga, 17 Apr 1999. CBF 8683, 6, Pantanal, Río Las Conchas, Estancia Las Conchas, arroyo “submontano” de “aguas blancas”, con alternancia de rápidos y pozas, 17° 33’ 58.3’’S 59° 28’ 17.1’’W, J. Sarmiento & K. Osinaga, 16 Apr 1999. MNKP 739 (589), 1, San Matías, Estación Cascabel, M.L. Argandoña, 8 Sep 1994. MNKP 742 (535), 2, San Matías, Estancia Cascabal, M.L. Argandoña, 8 Sep 1994. MNKP 3227 (4903), 2, Laguna Mandioré, K. Osinaga, 16 Jul 1998. MNKP 4727 (3613), Santo Corazon, río Santo Corazon, K. Osinaga & P. Coro, 28 Sep 1999. MNKP 4816 (6600), 3, río Santo Rosario, K. Osinaga & P. Coro, 1 Oct 1999. **Chiquitos:** MNKP 4877 (3355), 1, Carmen Rivero Torrez. río Otuquis, V. Fuentes & S. Sangueza, 1 Nov 1999. MNKP 4867 (4250), 1, río Tucavaca, V. Fuentes & S. Sangueza. MNKP 5232, 1, Laguna al lado derecho del Camino, 800 m antes de la localidad de Candelaria, M.E. Farell, M.Á. Velásquez & P. Hinojosa, 29 - 30 Dec 2003. MNKP 5328, 1, Localidad Candelaria, río Aguas Calientes, M.E. Farell, M.Á. Velásquez & G. Huanca, 2 Jul 2004. MNKP 5363, 3, Candelaria, río Tucavaca, M.E. Farell, M.Á. Velásquez & G. Huanca, 30 Jun 2004. **Germán Busch:** MNKP 3953 (3209), 4, poza en el camino a hacienda Santa Elena, K. Osinaga & J. Cardona, 31 Oct 1998.

###### Identification

Differs from *B. oenolaemus* by a longer pectoral fin (36-41 vs. 29-33 % of SL) and a shorter head (35-38 vs. 38-41 % of SL). Body (at least ventral portions of head and flanks) with a yellowish hue when alive, brachyostegal membrane without chromatophores. A brief redescription of *B. vittata* is provided by Kullander (1981). *Bujurquina vittata* is a small species reaching 7.9 cm SL (Hablützel, unpublished data).

### *Chaetobranchopsis australis* Eigenmann & Ward, 1907

*Chaetobranchopsis australe* Eigenmann & Ward, 1907

#### Type locality

Bahia Negra [Paraguay].

##### Bibliography for the study area

*Chaetobranchopsis australis*; Haseman (1911): locality (Puerto Suarez) and voucher material.

*Chaetobranchopsis australis*; Farell & Cancino (2007): locality (Río Tucavaca).

###### Examined material

**Bolivia: río Paraguay drainage: Santa Cruz: Ángel Sandoval:** CBF 8686, 1, Pantanal, Río Las Conchas, Estancia Las Conchas, arroyo “submontano” de “aguas blancas”, con alternancia de rápidos y pozas, 17° 33’ 58.3’’S 59° 28’ 17.1’’W, J. Sarmiento & K. Osinaga, 16 Apr 1999. MNKP 2459 (2175), 1, Hacienda Santa Helena, río Mercedes, R. Coca & Maraz, 22 Sep 1997. **Germán Busch:** MNKP 3929 (2997), 1, Puerto Suarez, Estación Verde, Laguna Cáceres, K. Osinaga & J. Cardona, 28 Oct 1998. MNKP 3947 (3033), 22, poza en el camino a la hacienda Santa Elena (Otuquis), K. Osinaga & J. Cardona, 31 Oct 1998.

###### Identification

*Chaetobranchopsis australis* belongs to the tribe Chaetobranchini, which are specialized filter feeder and have a large number of long gill rakers (around 50 or more on the first gill arch). Most other species of Neotropical cichlids have about 20 or less short gill rakers on the first gill arch. It is the only species of Chaetobranchini known to occur in the La Plata basin. *Chaetobranchopsis australis* is a medium-sized species reaching 12 cm SL (Kullander 2003).

### *Cichla piquiti* Kullander & Ferreira, 2006

*Cichla piquiti* Kullander & Ferreira, 2006

#### Type locality

Brazil: Pará: Rio Itacaiúnas at Caldeirão.

##### Bibliography for the study área

*Cichla piquiti*; Sarmiento *et al.* (2014): listing (Paraguay-Paraná Basin).

###### Examined material

None.

###### Identification

The genus *Cichla* can be readily distinguished from other South American cichlids by its large size, the deeply notched dorsal-fin margin, the small scales, the densely scaled unpaired fins and the conspicuous ocellus on the base of the caudal fin. *Cichla piquiti* and *C. kelberi* Kullander & Ferreira, 2006 are the only species of *Cichla* reported from the upper río Paraguay (both introduced). They can readily be distinguished on the number of vertical bands below the dorsal fin: three in *C. piquiti* vs. five in *C. kelberi*. *Cichla piquiti* is a large species reaching 43 cm SL (Kullander & Ferreira 2006).

### *Cichlasoma dimerus* (Heckel, 1840)

*Acara dimerus* Heckel, 1840

*Acara marginatus* Heckel, 1840

*Heros centralis* Holmberg, 1891

#### Type locality

Cujabá-Fluss.

##### Bibliography for the study area

*Aequidens portalegrensis*; Haseman (1911): localities (Puerto Suarez) and voucher material. *Cichlasoma dimerus*; Kullander (1983): locality (R. San Rafael, 4 km S Roboré) and voucher material.

*Cichlasoma boliviense*; Farell & Cancino (2007): locality (Río Roboré).

###### Examined material

**Bolivia: río Paraguay drainage: Santa Cruz: Angel Sandoval:** CBF 8623, 3, Pantanal, pantano con lagunilla, 17°04’10.5’’S 59°28’17.1’’W, J. Sarmiento & K. Osinaga, 19 Apr 1999. CBF 8651, 11, Pantanal, Lago Santa Elena, J. Sarmiento & K. Osinaga, 20-21 Apr 1999. MNKP 736 (537), 2, San Matías, Estancia Cascabel, M.L. Argandoña, 8 Sep 1994. MNKP 740 (535), 5, San Matías, Estancia Cascabal, M.L. Argandoña, 8 Sep 1994. MNKP 2416 (2112), 1, a 3 km de San Matías, debajo del puente, R. Coca & Maraz, 20 Sep 1997. MNKP 2418 (2156), 2, a 3 km de San Matías, debajo del puente, R. Coca, Maraz, 20 Sep 1997. MNKP 2426 (2193), 1, a 3 km de San Matías, debajo del puente, R. Coca & Maraz, 20 Sep 1997. MNKP 2441 (2209), 1, camino a curiche la Hormiga a 14 km de San Matías, Curichi Patujusal, R. Coca & Maraz, 21 Sep 1997. MNKP 3228 (6116), 7, Puerto Gonzalo, Laguna Mandioré, K. Osinaga, 16 Jul 1998. MNKP 3270 (4903), 4, Laguna Mandioré, K. Osinaga, 16 Jul 1998. MNKP 4746 (3518), 4, Curiche Tapera, K. Osinaga & P. Coro, 29 Sep 1999. MNKP 4612 (3453), 1, laguna, Hacienda Vista Hermosa, laguna, K. Osinaga & P. Coro, 21 Sep 1999. MNKP 6147, 1, río Bella Boca, V. Fuentes, 23 Nov 2000. **Chiquitos:** MNKP 4892 (3371), 1, río Bravo, V. Fuentes & S. Sangueza, 5 Nov 1999. MNKP 4850 (3410), 5, río Aguas Calientes, V. Fuentes, S. Sangueza, 31 Oct 1999. MNKP 3792, 1, río Esperancita, M. Velasquez & V. Fuentes, 2 Feb 2006. MNKP 5353, 2, río Aguas Calientes, zona El Playón, Aguas Calientes, M.E. Farell, M.Á. Velásquez & G. Huanca, 5 Jul 2004. **Germán Busch:** MNKP 1155 (1537), 1, Puerto Suarez, Laguna Cáceres, Palo Santo, K. Osinaga, R. Coca, V. Fuentes & A. Franco, 19 Feb 1996. MNKP 1156 (1548), 3, Puerto Suarez, Hacienda San Lorenzo, río San Lorenzo, K. Osinaga, R. Coca, V. Fuentes & A. Franco, 29 Feb 1996. MNKP 1180 (1667), 1, Puerto Suarez, Laguna Cáceres, K. Osinaga, R. Coca, V. Fuentes & A. Franco, 22 Feb 1999 (lot also contains 2 specimens of *Apistogramma borellii*). MNKP 1195 (1739), 1, curiche temporal, Laguna Cáceres, Puerto Suárez, K. Osinaga, R. Coca, V. Fuentes & A. Franco, 19 Feb 1996. MNKP 1218 (2261), 1, Puerto Quijarro, Tamarinero, Canal Tamengo, K. Osinaga, R. Coca, V. Fuentes & A. Franco, 26 Feb 1996. MNKP 1219 (2732), 10, Puerto Quijarro, bahia Cáceres, Canal Tamengo, K. Osinaga, R. Coca, V. Fuentes & A. Franco, 26 Feb 1996 (lot also contains 10 specimens of *Apistogramma borellii*). MNKP 1223 (4229), 2, Puerto Suarez, Laguna Cáceres, curiche temporal, K. Osinaga, R. Coca, V. Fuentes & A. Franco, 19 Feb 1996. MNKP 1327 (1561), 2, Puerto Suarez, Laguna Cáceres, Hacienda Arcoiris, K. Osinaga, V. Fuentes, A. Justiniano & E. Guzmán, 19 May 1996. MNKP 1394 (1491), 1, Puerto Suárez, Laguan Cáceres, P. Rebolledo, K. Osinaga, V. Fuentes, L. Paniagua & L. Paredes, 9 Aug 1996. MNKP 1427 (1528), 1, Puerto Suarez, Laguna Cáceres, Castrillo, P. Rebolledo, K. Osinaga, V. Fuentes, L. Paniagua & L. Paredes, 9 Aug 1996. MNKP 1429 (1531), 7, Puerto Suarez, Laguna Cáceres, Hacienda Arcoiris, P. Rebolledo, K. Osinaga, V. Fuentes, L. Paniagua & L. Paredes, 4 Aug 1996. MNKP 1516 (2255), 1, Puerto Quijarro, Tamarinero, Laguna Cáceres, P. Rebolledo, K. Osinaga, L. Paniagua, L. Paredes & V. Fuentes, 13 Aug 1996. MNKP 2193 (2035), 44, El Carmen, zona de inundación, estancia Campo el Medio, P. Rebolledo, R. Coca, V. Fuentes & G. Soto, May 1997. MNKP 2197 (2051), 1, El Carmen, zona de inundación, estancia Campo el Medio, P. Rebolledo, R. Coca, V. Fuentes & G. Soto, May 1997. MNKP 2293 (2046), 1, Puerto Busch, Isla Santa Rosa, río Paraguay, R. Coca, V. Fuentes & G. Soto, 14 May 1997. MNKP 3902 (3017), 7, Puerto Suarez, río Pimiento, K. Osinaga & J. Cardona, 27 Oct 1998. MNKP 3907 (3201), 2, Puerto Suarez, Laguna Cáceres, Pto. Arcoiris, río Pimiento, K. Osinaga, J. Cardona, 27 Oct 1998. MNKP 3928 (2995), 2, Puerto Suarez, Laguna Cáceres, Est. Verde, K. Osinaga & J. Cardona, 28 Oct 1998. MNKP 3960 (4321), 33, Poza en el camino a Hacienda Santa Elena, río Otuquis drainage, K. Osinaga & J. Cardona, 31 Oct 1998. MNKP 4021 (3018 and 3026), 2, El Carmen, zona de inundanción, Estancia Campo el Medio, K. Osinaga & J. Cardona, 9 Nov 1998. MNKP 4037 (3037), 19, El Carmen, Estancia Campo en Medio, curichi a 2 km suroeste, K. Osinaga & J. Cardona, 10 Nov 1998. MNKP 4046 (3261), 18, El Carmen, Estancia Campo en Medio, curichi a 2 km suroeste, K. Osinag &, J. Cardona, 10 Nov 1998. MNKP 4319, 2, Santa Ana, río la Canoa, V. Fuentes, 21 Oct 2000. MNKP 5085 (4327), 2, Sta. Ana, Poza La Banda, Santa Ana, V. Fuentes, 20 Oct 2000. **río Bermejo drainage: Tarija: Arce:** CBF 8605, 4, Río Bermejo, Ciudad de Bermejo: Punto 2, 22°47’’S 64°20’W, J. Horton, 27 Dec 1993.

###### Identification

As described in Kullander (1983). Most adult or sub-adult specimen of *Cichlasoma boliviense*, a species from adjacent areas in the Amazon, have scale-base spots on the posterioventral parts of the sides, resulting in a mottled pattern, which are absent in *C. dimerus*. The two species overlap in measurements and meristics (Kullander 1983), making it virtually impossible to distinguish them by morphology in some cases. However, no specimen with mottled colouration pattern have yet been observed in the La Plata drainage, indicating that *C. boliviense* is indeed restricted to the Amazon. *Cichlasoma dimerus* is a medium-sized species reaching 11.7 cm SL (Kullander 1983).

### *Crenicichla lepidota* Heckel, 1840

*Crenicichla lepidota* Heckel, 1840

*Crenicichla edithae* Ploeg, 1991

#### Type locality

Rio-Guaporè.

##### Bibliography for the study area

*Crenicichla lepidota*; Haseman (1911): locality (Puerto Suarez) and voucher material.

*Crenicichla lepidota*; Terrazas Urquidi (1970): listing (R. Bermejo and R. Paraguay).

*Crenicichla lepidota*; Farell & Cancino (2007): localities (Río Otuquis drainage).

*Crenicichla lepidota*; Varella (2011): bibliography.

###### Examined material

**Bolivia: río Paraguay drainage: Santa Cruz: Angel Sandoval:** CBF 8652, 1, Pantanal, Lago Santa Elena, Paraguay drainage, J. Sarmiento & K. Osinaga, 20-21 Apr 1999. CBF 8680, 1, Pantanal, Río Las Conchas, Estancia Las Conchas, arroyo “submontano” de “aguas negras”, con presencia de grandes pozas, rápidos y cataratas, 17° 33’ 58.3’’S 59° 28’ 17.1’’W, Sarmiento & K. Osinaga, 17 Apr 1999. MNKP 2417 (2113), 2, río a 3 km de San Matías, debajo del puente, R. Coca & Maraz, 20 Sep 1997. MNKP 2420 (2158), 10, río a 3 km de San Matías, debajo del puente, R. Coca & Maraz, 20 Sep 1997. MNKP 2425 (2187), 1, río a 3 km de San Matías, debajo del puente, R. Coca & Maraz, 20 Sep 1997. MNKP 2435 (2169), 2, Curichi Patujusal, camino a la curiche La Hormiga, 14 km a San Matías, R. Coca & Maraz, 21 Sep 1997. MNKP 4716 (3490), 1, Santo Corazón, río Santo Corazón, K. Osinaga & P. Coro, 28 Sep 1999. MNKP 4747 (3519), 2, Curiche Tapera, K. Osinaga & P. Coro, 29 Sep 1999. MNKP 4105 (3611), 1, pantano, viejo cauce de río, J. Sarmiento, K. Osinaga, F. Osorio & R. Bueno, 23 Apr 1999. MNKP 4587 (3506), 1, Area de Manejo Integral San Matías, laguna, Hacienda Vista Hermosa, Osinaga & P. Coro, 20 Sep 1999. MNKP 3139 (3924), 2, río Pando, Puerto Gonzalo & K. Osinaga, 11 Julio 1998. MNKP 3223 (4151), 1, Laguna Uberaba, K. Osinaga, 4 Jul 1998. MNKP 4105 (3641), 6, Área de Manejo Integral San Matías, Hacienda Pantano Caribe, río Caribe, K. Osinaga & P. Coro, 24 Sep 1999. MNKP 4680 (3643), 4, Area de Manejo Integral San Matías, Hacienda Pantano Caribe, río Caribe, K. Osinaga & P. Coro, 27 Sep 1999. MNKP 4570 (3664), 4, laguna, Hacienda Vista Hermosa, K. Osinaga & P. Coro, 19 Sep 1999. MNKP 4645 (3870), 5, Area de Manejo Integral San Matías, Hacienda Pantano Caribe, río Caribe, K. Osinaga & P. Coro, 27 Sep 1999. MNKP 3223 (4187), 1, Laguna Mandioré, K. Osinaga, 15 Jul 1998. **Chiquitos:** MNKP 4762 (3622), 7, Aguas Calientes, río Aguas Calientes, K. Osinaga & P. Coro, 26-27 Sep 1999. MNKP 4864 (4123), 2, ?, V. Fuentes & S. Sangueza, Nov 1999. MNKP 5256, 1, laguna al lado del camino, 800 m antes de llegar a Candelaria, M.E. Farell, M.Á. Velásquez & P. Hinojosa, 30 Dec 2003. MNKP 5269, 31, Candelaria, río Tucavaca, M.E. Farell, M.Á. Velásquez & P. Hinojosa, 29 Dec 2003 (lot also contains 1 specimen of *C. semifasciata*). MNKP 5296, 1, Candelaria, río Aguas Calientes, M.E. Farell, M.Á. Velásquez & G. Huanca, 2 Jul 2004. MNKP 5305, 5, laguna al lado derecho del camino, 800 m antes de llegar a la localidad de Candelaria, M.E. Farell, M.Á. Velásquez & G. Huanca, 1 Jul 2004. **Germán Busch:** MNKP 1182 (1670), 1, Puerto Suarez, Palo Santo, Laguna Cáceres, K. Osinaga, R. Coca, V. Fuentes & A. Franco, 21 Feb 1996. MNKP 1403 (1498), 5, Puerto Quijarro, Tamarinero, Laguna Cáceres, P. Rebolledo, K. Osinaga, V. Fuentes, L. Paniagua & L. Paredes, 15 Aug 1996. MNKP 1404 (1499), 1, Puerto Quijarro, Tamarinero, Laguna Cáceres, P. Rebolledo, K. Osinaga, V. Fuentes, L. Paniagua & L. Paredes, 13 Aug 1996. MNKP 1422 (1521), 1, Puerto Quijarro, Tamarinero, Laguna Cáceres, P. Rebolledo, K. Osinaga, V. Fuentes, L. Paniagua & L. Paredes, 8 Aug 1996. MNKP 1424 (1524), 1, Puerto Suarez, Laguna Cáceres, zona de la Bamba antigua, P. Rebolledo, K. Osinaga, V. Fuentes, L. Paniagua & L. Paredes, 11 Aug 1996. MNKP 1484 (1729), 1, Puerto Suarez, Laguna Cáceres, P. Rebolledo, K. Osinaga, L. Paniagua, L. Paredes & V. Fuentes, 9 Aug 1996. MNKP 1425 (1525), 4, Puerto Quijarro, Tamarinero, Laguna Cáceres, P. Rebolledo, K. Osinaga, V. Fuentes, L. Paniagua & L. Paredes, 13 Aug 1996. MNKP 1526, 5, Puerto Quijarro, Laguna Cáceres, P. Rebolledo, K. Osinaga, V. Fuentes, L. Paniagua, L. Paredes, 13 de Ago 1996. MNKP 2206 (2083), 3, El Carmen, zona de inundación, Estancia Campo el Medio, P. Rebolledo, R. Coca, V. Fuentes & G. Soto, May 1997. MNKP 3945 (3011), 6, poza en el camino Hacienda Santa Elena, K. Osinaga & J. Cardona, 31 Oct 1998. MNKP 3973 (3036), 4, Camino a Hacienda Santa Elena, K. Osinaga & J. Cardona, 1 Nov 1998. MNKP 3900 (2987), 2, Puerto Suarez, río Jordano Pimiento, K. Osinaga & J. Cardona, 27 Oct 1998. MNKP 3993 (3000), 4, Otuquis, Bahia Corea, K. Osinaga, J. Cardona, A. Justiniano & M. Chavez, 5 Nov 1998.

###### Identification

A redescription of *C. lepidota* is provided by Kullander (1982) and Varella (2011). It differs from *C. semifasciata* by a less depressed head and a longer and more slender snout and from *C. vittata* by lower longitudinal scale count (34-51 vs. about 79-90 E1 scales). *Crenicichla ploegi* and *C. semicincta* are similar and parapatric species. The form can be distinguished from *C. lepidota* by a higher scale count (58-71 E1 scales) and the colour pattern (Varella *et al.* 2018). The latter differs from *C. lepidota* by lacking a ring of iridescent scales around the lateral spot and having this spot elongated and nearly integrated in the midlateral longitudinal band. *Crenicichla semicincta*, however, has not been reported from the La Plata system and appears to be restricted to the upper río Madera drainage. The species is medium-sized reaching 18.9 mm SL (Varella 2011).

### *Crenicichla semifasciata* (Heckel, 1840)

*Batrachops semifascaitus* Heckel, 1840

*Acharnes chacoensis* Holmberg, 1891

*Boggiania ocellata* Perugia, 1897

*Crenicichla simoni* Haseman, 1911

#### Type locality

Flusse Paragauy bei Caiçara.

##### Bibliography for the study area

*Crenicichla simoni*; Haseman (1911): locality (Puerto Suarez) and voucher material.

*Batrachops ocellatus*; Terrazas Urquidi (1970): listing (R. Paraguay). *Batrachops semifasciatus*; Terrazas Urquidi (1970): listing (R. Paraguay). *Crenicichla semifasciata*; Farell & Cancino (2007): locality (Río Tucavaca).

###### Examined material

**Bolivia: río Paraguay drainage: Santa Cruz: Chiquitos:** MNKP 5269, 1, Candelaria, río Tucavaca, M.E. Farell, M.Á. Velásquez & P. Hinojosa, 29 Dec 2003 (lot also contains 31 specimens of *C. lepidota*). **Germán Busch:** MNKP 1216 (2259), 1, Puerto Quijarro, Tamarinero, Canal Tamengo, K. Osinaga, R. Coca, V. Fuentes & A. Franco, 26 Feb 1996. MNKP 1220 (2733), 1, Puerto Suarez, Laguna Cáceres, Palo Santo, K. Osinaga, R. Coca, V. Fuentes & A. Franco, 23 Feb 1996.

###### Identification

Differs from all other *Crenicichla* species in the Paraguay-Paraná drainage by a wider head and a shorter and broader snout. In terms of coloration it differs from the other species by having the base of the scales pigmented, giving the sides a finely reticulated appearance. *Crenicichla semifasciata* is a medium-sized species reaching 18.5 cm SL (Varella 2011).

### *Crenicichla vittata* Heckel, 1840

*Crenicichla vittata* Heckel, 1840

#### Type locality

Flusse Cuyaba; … Flusse Paraguay

##### Bibliography for the study area

*Crenicichla vittata*; Terrazas Urquidi (1970): listing (R. Paraguay).

###### Examined material

**Bolivia: río Paraguay drainage: Santa Cruz: Chiquitos:** MNKP 4853 (3348) río Aguas Calientes, V. Fuentes & S. Sangueza, 1 Nov 1999. **Germán Busch:** MNKP 1130 (1384), 1, Puerto Suarez, San Eugenio, Laguna Cáceres, K. Osinaga, R. Coca, V. Fuentes & A. Franco, 19 Feb 1996.

###### Identification

*Crenicichla vittata* is straightforward to identify as it is the only *Crenicichla* species in the Upper Río Paraguay basin with about 83-95 scales in the longitudinal row (Kullander 1981; Ploeg 1991). Other species of *Crenicichla* known from other drainage systems with a similar scale count lack a suborbital stripe (Ploeg 1991). Differs from *C. semifasciata* by a less depressed head and a longer and slenderer snout and from *C. lepidota* by higher longitudinal scale count (about 83-95 vs. about 34-45). It is a large species reaching a size of 29.4 cm SL (Varella 2011). See figure 6 for general appearance. A brief redescription of *C. vittata* is provided by Kullander (1981).

**Fig. 6:**
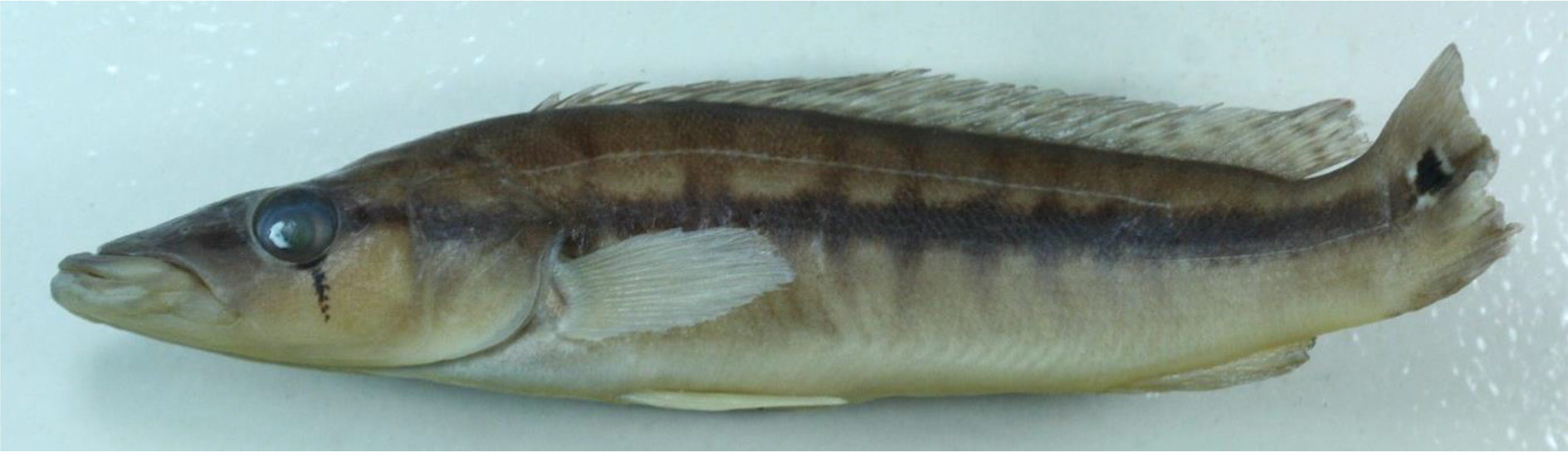
*Crenicichla vittata* from the río Aguas Calientes, río Paraguay drainage, Chiquitos, Santa Cruz, Bolivia (MNKP 3348; tail is bent to the left).

### *Crenicichla* sp. (unidentified juveniles)

#### Examined material

**Bolivia: río Paraguay drainage: Santa Cruz: Ángel Sandoval:** MNKP 4780 (3570), many juveniles, Bahia Jesus.

### *Gymnogeophagus balzanii* **(Perugia, 1891)**

*Geophagus balzanii* Perugia, 1891

*Geophagus duodecimspinosus* Boulenger, 1895

*Gymnogeophagus cyanopterus* Miranda Ribeiro, 1918

#### Type locality

Villa Maria (Matto Grosso), Rio Paraguay a 15°.

##### Bibliography for the study area

*Geophagus balzanii*; Haseman (1911): locality (Puerto Suarez) and voucher material.

*Gymnogeophagus balzanii*; Farell & Cancino (2007): locality (Río Tucavaca).

###### Examined material

**Bolivia: río Paraguay drainage: Santa Cruz: Ángel Sandoval:** MNKP 4797 (3522), 4, Santo Rosario, río Santo Rosario, K. Osinaga & P. Coro, 1 Oct 1999. **Chiquitos:** MNKP 5313, 1, Candelaria, río Tucavaca, M.E. Farell, M.Á. Velásquez & G. Huanca, 30 Jun 2004.

###### Identification

*Gymnogeophagus balzanii* differs from similar species in the study area by having the dorsal fin scaled and lacking a caudal spot. Males develop a conspicuous nuptial bulb on the forehead during the reproductive season. It is the only species of *Gymnogeophagus* known to occur in the upper reaches of the río Paraguay drainage. It is a medium-sized species reaching 12 cm SL (Kullander 2003).

### *Laetacara dorsigera* (Heckel, 1840)

*Acara dorsiger* Heckel, 1840

#### Type locality

Sümpfe in der Nähe des Paraguay-Flusses bei Villa Maria.

##### Bibliography for the study area

*Acara dorsigera*; Haseman (1911): locality (Puerto Suarez) and voucher material.

###### Examined material

**Bolivia: río Paraguay drainage: Santa Cruz: Ángel Sandoval:** CBF 8630, 48, Pantanal, pantano con lagunilla, 17° 04’ 10.5’’S 59° 28’ 17.1’’W, J. Sarmiento & K. Osinaga, 19 Apr 1999. CBF 8650, 16, Pantanal, Lago Santa Elena, J. Sarmiento & K. Osinaga, 120-21 Apr 1999. MNKP 735 (589), 2, San Matías, Estación Cascabel, M.L. Argandoña, 8 Sep 1994. MNKP 741 (535), 3, San Matías, Estancia Cascabal, M.L. Argandoña, 8 Sep 1994. MNKP 2422 (2160), 2, San Matías debajo del puente, R. Coca & Maraz, 20 Sep 1997. MNKP 3229 (6116), 2, Puerto Gonzalo, Laguna Mandioré, K. Osinaga, 16 Jul 1998. MNKP 4623 (3453), 1, laguna, Hacienda Vista Hermosa, laguna, K. Osinaga & P. Coro, 21 Sep 1999.

###### Identification

As described in Ottoni & Costa (2009). *Laetacara* species can be distinguished from similar cichlid genera by the presence of scales on the preoperculum (Kullander 1986). *Laetacara dorsigera* is a small species reaching 4.0 cm SL (Ottoni 2018).

### *Mesonauta festivus* (Heckel, 1840)

*Heros festivus* Heckel, 1840

#### Type locality

Fluss Guaporè und dessen nahe gelegenen Moräste.

##### Bibliography for the study area

###### Examined material

**Bolivia: río Paraguay drainage: Santa Cruz: Ángel Sandoval:** MNKP 2414 (1984, 2 and 2155, 1), 3, San Matías a 3 km bajo el puente, R. Coca & Maraz, 20 Sep 1997. MNKP 3514 (4547), 1, Area de Manejo Integrado San Matías, Bahia Jesús, K. Osinaga & P: Coro, 30 Sep 1999. MNKP 3254 (3933), 3, Laguna Mandioré, K. Osinaga, 16 Jul 1998. MNKP 5204 (4689), 3, Laguna La Selva, V. Fuentes, 17 Dec 2000.

###### Identification

The genus *Mesonauta* can easily be distinguished from other cichlid genera by the presence of a conspicuous black stripe from the lower jaw to the rayed part of the dorsal fin. Further characteristics are the disciform body, the long pelvic spine (reaching beyond the origin of the anal fin) and the high number of anal fin spines (VII-XI; Kullander & Silfvergrip 1991). It is a small species reaching 8 cm SL (Kullander & Silfvergrip 1991). No other species of *Mesonauta* have been reported from the Río Paraguay basin. See Figure 7 for general appearance of the species.

**Fig. 7:**
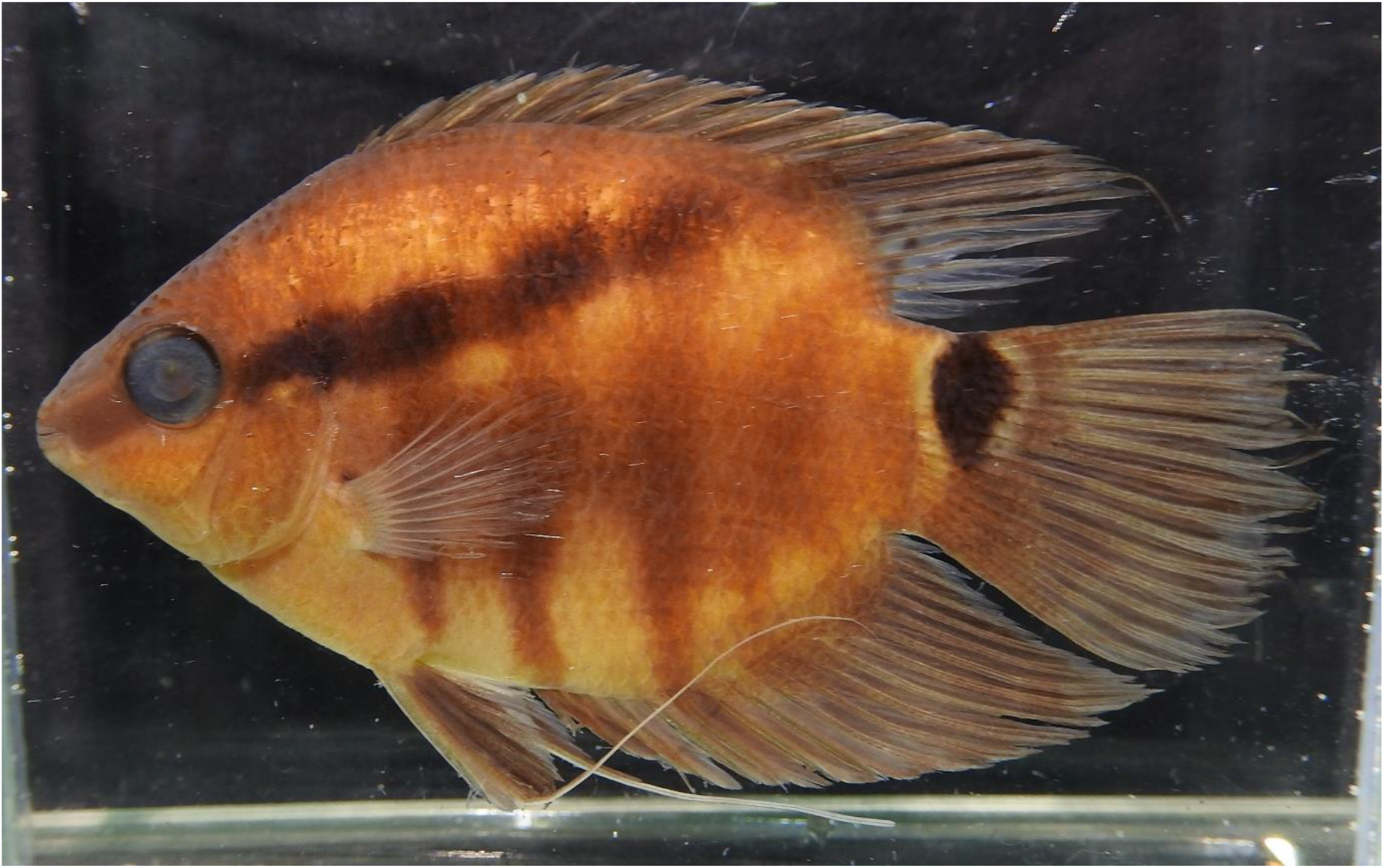
*Mesonauta festivus* from San Matias, debajo del puente, río Paragauy drainage, Angel Sandoval, Santa Cruz, Bolivia (MNKP 2414 (1984 or 2155)).

### *Satanoperca pappaterra* (Heckel, 1840)

*Geophagus pappaterra* Heckel, 1840

#### Type locality

Rio-Guapore.

##### Bibliography for the study area

###### Examined material

**Bolivia: río Paraguay drainage: Santa Cruz: Ángel Sandoval:** MNKP 725 (532), 1, San Matías, Estancia Cascabel, M.L. Argandoña, 8 Sep 1994. **Germán Busch:** MNKP 5203 (4147), 1, Laguna La Selva, V. Fuentes, 17 Dec 2000.

###### Identification

No unambiguous diagnostic characters are currently known for *Satanoperca pappaterra* to separate it from *S. jurupari*. However, the species has frequently been reported from the study area and is the only species of its genus occurring in the La Plata basin. From other geophagine cichlid genera, *Satanoperca* can be distinguished by the absence of scales on the dorsal fin and the lack of a suborbital stripe. *Satanoperca pappaterra* is a medium-sized species reaching 17.4 cm SL (Kullander 2003). See Figure 8 for general appearance of the species.

**Fig. 8:**
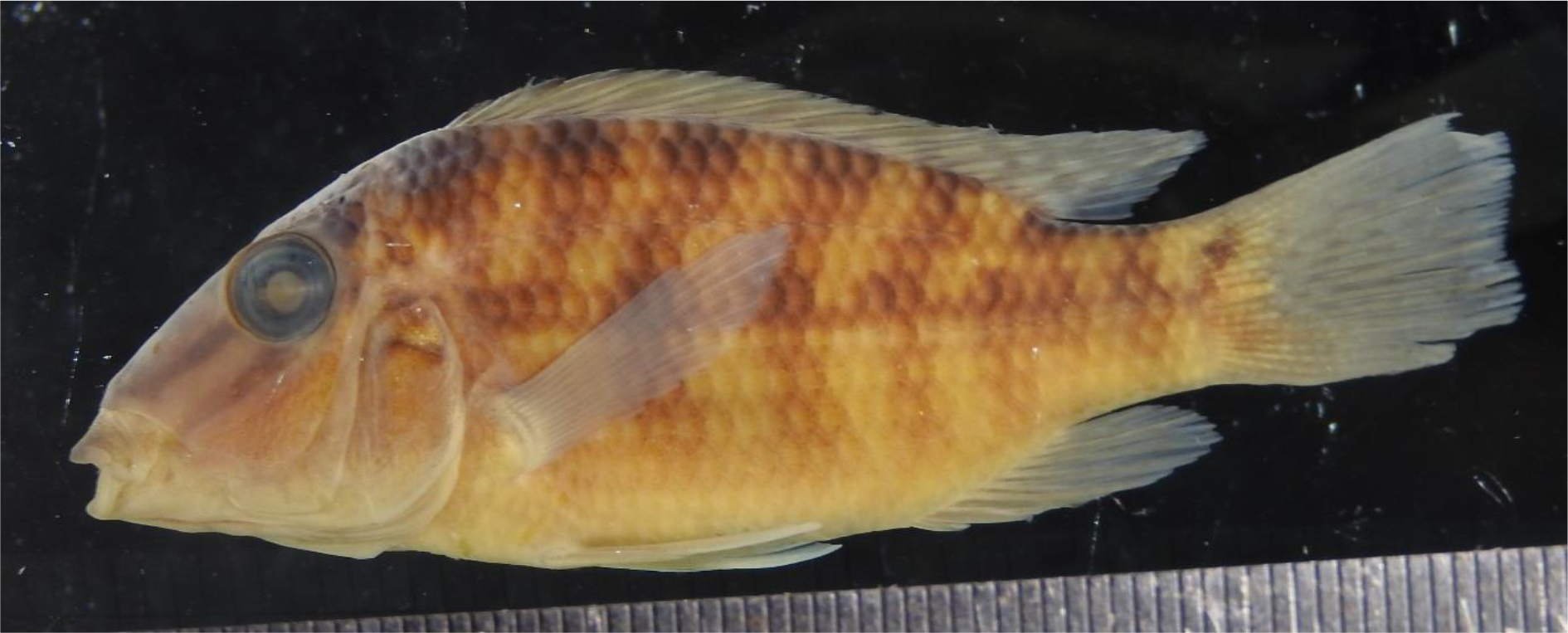
*Satanoperca pappaterra* from Laguna La Selva, río Paraguay drainage, Germán Busch, Santa Cruz, Bolivia (MNKP 5203).

**Key to species of the La Plata drainage in Bolivia and the upper río Paraguay in Brazil:**

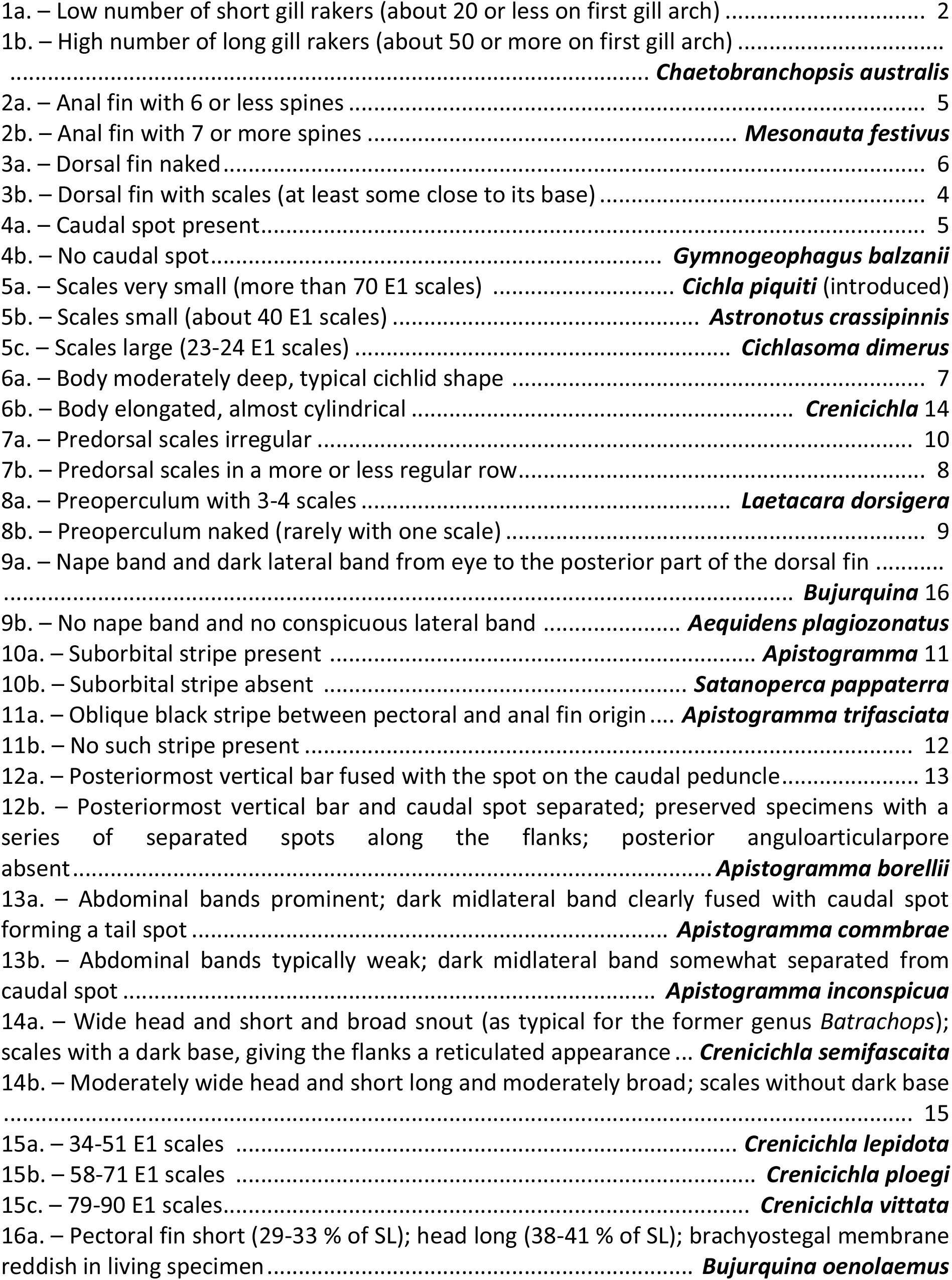
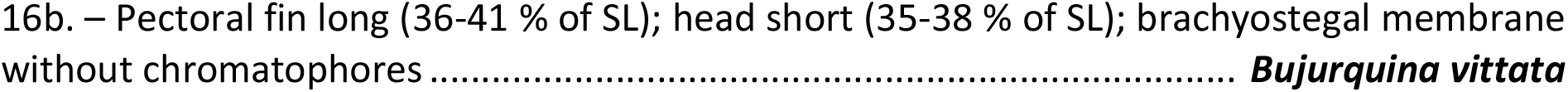

## Discussion

The examination of 749 specimens in 162 museum lots revealed 16 species of cichlid fishes. Virtually all records stem from the upper río Paraguay and only one lot (*Cichlasoma dimerus*) has been collected in the río Bermejo drainage. We can also report only one literature report from the Bermejo river (Chaetobranchus sp.; Sarmiento *et al.* (2019)) and río Pilcomayo (*Bujurquina vittata*; Terrazas Urquidi (1970).

### Species considered not to occur in the La Plata drainage of Bolivia

Sarmiento & Barrera (1997) and Sarmiento *et al.* (2019) reported *Acaronia* sp. from the río Pilcomayo basin. According to the CBF catalog, these specimens are from the río Bermejo. These samples could not be located in the collection and the occurrence of this Amazonian species so far south as the río Bermejo is highly questionable.

Terrazas Urquidi (1970) listed *Apistogramma taeniatum* (= *A. taeniata*) and *Aequidens tetramerus* for the Bolivian part of the río Paraguay drainage, but did not provide any further information about specimen, preventing verification. The former species is restricted to the lower rio Tapajós drainage in Brazil (Kullander 2003) and its report is certainly based on a misidentification. *Aequidens tetramerus s.l.* occurs in the upper río Madera (Kullander 1986; Hablützel & Pantel 2017), but has not been reported from the La Plata drainage elsewhere, making this report dubious.

While the occurrence of *Chaetobranchopsis australis* is certain for the río Paraguay drainage in Bolivia, the report of *Chaetobranchopsis* sp. from the río Bermejo drainage (Sarmiento *et al.* 2019) appears odd. We have no knowledge of the possible occurrence of this species from the Argentinian section of the río Bermejo and it seems that the species is restricted to the río Paraguay drainage (possibly including the río Pilcomayo). We have not seen any specimens from the río Bermejo and think this report should be considered with caution.

### Identification issues

Discrimination between *Cichlasoma boliviensis* and *Cichlasoma dimerus* was based on sampling location, rather than diagnostic morphological traits. Subadults and adults of the former species typically (but not always) show a specific pattern of spots on the posterioventral portion of the flanks, giving them a mottled appearance. This colouration has never been reported from the La Plata basin, even though a large number of specimens have been screened. It appears reasonable to assume that the two species are allopatric and the watershed corresponds to the distribution boundary.

Juveniles that do not show the diagnostic characters of their respective species were only identified at genus level. This applies in particular to species of the genus *Apistogramma*, which might be impossible to distinguish morphologically at juvenile stages.

### The taxonomic status of *Crenicichla ocellata*

*Crenicichla ocellata* (Perugia, 1897) has been reported for the Bolivian part of the La Plata basin by Terrazas Urquidi (1970). This name, has been considered a synonym of *C. semifasciata* by several taxonomists (Ploeg 1991; Kullander 2003; Varella 2011). The Catalog of Fishes (Fricke *et al.* 2018), however, lists *C. ocellata* as a valid species with reference to several recent taxonomic studies. These publications, however, do not treat *C. ocellata* in the main text, but merely list the holotype among the examined material (e.g. Piálek *et al.* 2010, 2015; Casciotta *et al.* 2013; Říčan *et al.* 2017). In one publication, these authors list *C. ocellata* unter its original name *Boggiania ocellata* (Piálek *et al.* 2010), indicating that the authors did not intend to recognize *B. ocellata* as a distinct species, but used its original name to indicate the holotype. However, other researchers subsequently started to list *C. ocellata* as a valid species (Koerber *et al.* 2017). Because no argument for or against the synonymy of *B. ocellata* in *C. semifasciata* is provided in recent taxonomic studies, we follow the authors that treat this taxonomic question explicitly or implicitly (Ploeg 1991; Kullander 2003; Varella 2011) and continue to consider *B. ocellata* as a synonym of *C. semifasciata*.

### Completeness of the list

Most of the cichlid species that have been reported from the La Plata drainage in adjacent areas in Argentina, Brazil and Paraguay could be confirmed for Bolivia. One notable exception is *Apistogramma inconspicua*, which is, within the río Paraguay drainage, still only known from three specimen collected by Haseman in 1909 at a single location in Brazil (Est. Mato Grosso, R. Paraguay system, Cáceres; Kullander, 1982c). At least one of these specimens has, however, the lateral band continuous with the tail spot, which is characteristic for *A. commbrae*. A more detailed examination of these specimens is therefore necessary to confirm the presence of *A. inconspicua* in the río Paraguay drainage.

The introduced Amazonian *Cichla piquiti* has first been reported from the río Paraguay drainage in Bolivia by Sarmiento *et al.* (2014). Marques & Resende (2005) reported on *Cichla* cf. *monoculus* just north of Corumbá, a Brazilian city close to the Bolivian border. According to Kullander & Ferreira (2006) this introduced population belongs to *Cichla piquiti*. The occurrence of this species in Bolivia therefore appears possible, but unfortunately, no Bolivian specimens were accessible for examination.

Survey of the fish species in the Brazilian part of the Pantanal resulted in a cichlid species list that is almost identical to the one presented here (Menezes *et al.* 2000; Marcelino Polaz *et al.* 2014). Only *Bujurquina oenolaemus*, which is endemic to the río Aguas Calientes in Bolivia, was not recorded from Brazil. Menezes *et al.* (2000) and Varella (2011) reported on additional undescribed species of *Crenicichla*, whose distribution, however, may be restricted to the headwaters of the río Paraguay in Brazil. In 2018, one species was formally described as *Crenicichla ploegi* Varella, Loeb, Lima & Kullander, 2018 (Varella *et al.* 2018).

Our rarefaction analysis shows that the curve is flattening out (Figure 9), indicating that a large sampling effort has to be conducted to find additional species. We conclude that the species lists for the Bolivian La Plata basin is complete or at least almost complete and provides a sound basis for identification purposes, biodiversity inventories and ecological studies.

**Fig. 9:**
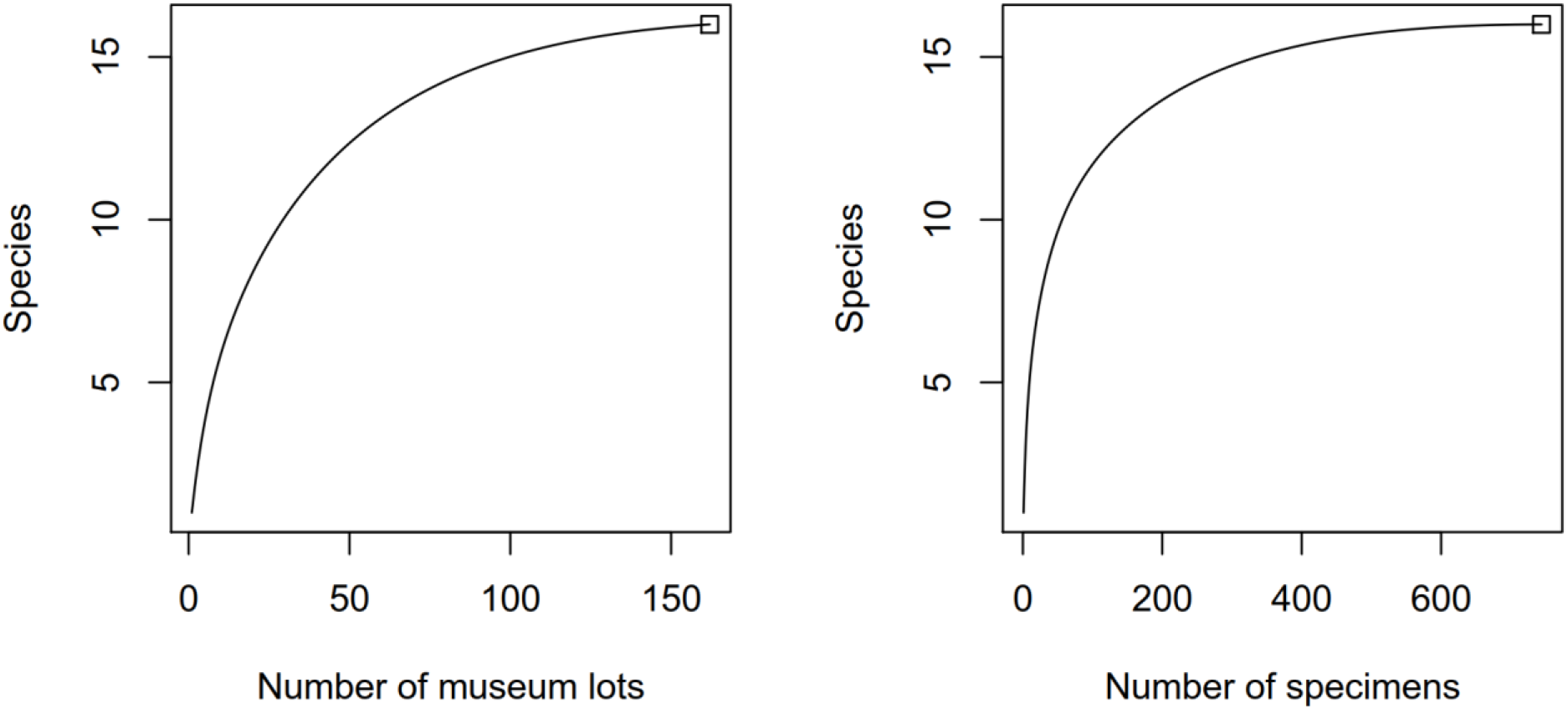
Rarefaction curves to visualize completeness of species sampling. On the left, museum lots were considered as individual species reports, while on the right the analysis was repeated at the level of individual voucher specimens.

## Conclusions

We report the occurrence of 17 species of cichlids in the Bolivian part of the La Plata drainage. Of those, 16 are documented with voucher specimens. The presence of the 17^th^ species (Cichla piquiti) appears reliably due to multiple reports of this species in neighbouring areas in Brazil. Our rarefaction analysis supports the notion that sampling was exhaustive and that this species list is complete or at least almost complete. The diagnostic characters mentioned in the text and the dichotomous key should enable reliable identification of cichlids in the field.

## Acknowledgements

We wish to thank Karina Osinaga and Kathia Riveiro (MNK, Santa Cruz, Bolivia), and Soraya Barrera and Jaime Sarmiento (CBF, Bolivia) for their hospitality during research visits and granting access to their collections. This research was financially supported by grants to PIH from the Deutsche Cichliden-Gesellschaft (DCG) and the Swiss Academy of Natural Sciences.

